# Cryo-electron tomographic investigation of native hippocampal glutamatergic synapses

**DOI:** 10.1101/2024.04.01.587595

**Authors:** Aya Matsui, Cathy J. Spangler, Johannes Elferich, Momoko Shiozaki, Nikki Jean, Xiaowei Zhao, Maozhen Qin, Haining Zhong, Zhiheng Yu, Eric Gouaux

**Author notes:** These authors contributed equally to this work (Aya Matsui, Cathy J. Spangler). Correspondence to Eric Gouaux.

## Abstract

Chemical synapses are the major sites of communication between neurons in the nervous system and mediate either excitatory or inhibitory signaling [1]. At excitatory synapses, glutamate is the primary neurotransmitter and upon release from presynaptic vesicles, is detected by postsynaptic glutamate receptors, which include ionotropic AMPA and NMDA receptors [2, 3]. Here we have developed methods to identify glutamatergic synapses in brain tissue slices, label AMPA receptors with small gold nanoparticles (AuNPs), and prepare lamella for cryo-electron tomography studies. The targeted imaging of glutamatergic synapses in the lamella is facilitated by fluorescent pre- and postsynaptic signatures, and the subsequent tomograms allow for identification of key features of chemical synapses, including synaptic vesicles, the synaptic cleft and AuNP-labeled AMPA receptors. These methods pave the way for imaging brain regions at high resolution, using unstained, unfixed samples preserved under near-native conditions.

## Main text

Chemical synapses underpin the majority of communication sites between neurons and are comprised of a presynaptic terminal, a synaptic cleft and a postsynaptic apparatus [1]. Signal transduction at synapses involves voltage-dependent and calcium-stimulated release of neurotransmitter from presynaptic vesicles, typically raising the concentration of neurotransmitter in the cleft as much as 10^5^-fold [4]. The bolus of neurotransmitter in turn promotes activation of synaptic and perisynaptic receptors, which are primarily located on postsynaptic specializations, thereby inducing depolarization, calcium entry into dendrites, and propagation of the neuronal impulse. At glutamatergic synapses, the neurotransmitter signal is quenched by clearance from synaptic and extracellular spaces by binding to and subsequent uptake via sodium-coupled transporters. Despite the central roles that chemical synapses play in the development and function of the nervous system, and their association with neurological disease and dysfunction, the molecular structure of the synapse has remained elusive, largely due to challenges associated with imaging at high resolution.

The first images of chemical synapses were derived from electron microscopy studies of fixed and stained neuronal and neuromuscular synapses [5]. Chemical fixation, dehydration and heavy metal staining during the sample preparation process, in combination with early electron microscopes and recording methods, severely constrained the resolution of the images and likely distorted molecular features. Nevertheless, it was possible to make out presynaptic vesicles, the synaptic cleft, and the postsynaptic density. Subsequent electron microscopy studies, using both cultured neurons and brain tissue-derived samples, together with imaging studies of biochemically isolated synaptosomes, have provided progressively higher resolution visualizations of synapses and their surrounding structures [6–8]. More recently, electron microscopy experiments, together with light microscopy studies, suggest that in excitatory, glutamatergic synapses, there is a transsynaptic organization of presynaptic and postsynaptic components that aligns sites of transmitter release and detection [9–13].

Here we report approaches for imaging glutamatergic synapses from the CA1 region of the hippocampus, a well-studied region of the brain that plays crucial roles in memory and learning and that includes the pyramidal cell, CA3-CA1 Schaffer collateral synapses. To enable identification and visualization of glutamatergic synapses, we developed several key methods. First, we generated an engineered mouse line to pin-point excitatory synapses via fluorescence methods. Second, we developed a GluA2-specific antibody fragment [14] gold nanoparticle (AuNP) conjugate [15] to localize AMPA subtype glutamate receptors. Third, we optimized sample preparation from unfixed hippocampal brain tissue, together with the application of cryo-protectants and high pressure freezing (HPF). With the frozen slices in hand, we next employed cryo-focused ion beam, scanning electron microscopy (cryo-FIB/SEM) milling methods [16–18] to produce thin lamella, in turn using pre- and postsynaptic fluorescence signals to guide imaging by cryo-electron tomography (cryo-ET). By combining all of these approaches, we created an experimental workflow that allows us to image excitatory synapses from unfixed, chemically unstained brain tissue by cryo-ET at near nanometer resolution, thereby opening the door to the direct localization of molecules involved in synapse structure, organization and function.

### Anti-GluA2 15F1 Fab conjugated to a single gold nanoparticle

To provide an electron-dense marker for labeling individual, GluA2-containing AMPARs at glutamatergic synapses in cryo-electron tomograms, we developed a strategy to label the anti-GluA2, 15F1 Fab [14, 17] with a single gold nanoparticle (AuNP) based on previously determined methods to label antibody fragments with AuNPs [19, 20]. To accomplish this, we synthesized uniform ∼3 nm 3-mercaptobenzoic acid (3-MBA) thiolate-protected gold nanoparticles by established methods [15]. AuNPs were coupled to an anti-GluA2 15F1 Fab engineered with an extended heavy chain containing a single free cysteine, thus enabling a thiol exchange reaction with 3-MBA and covalent coupling of the Fab to the AuNP (Figure 1A, Supplemental Figure 1). By optimization of the relative ratio of Fab and AuNP, we maximized the formation of a 1:1 Fab:AuNP species (Figure 1B). Following Fab conjugation, AuNPs were PEGylated to minimize aggregation of the nanoparticles (Supplemental Figure 1).

**Figure 1.**
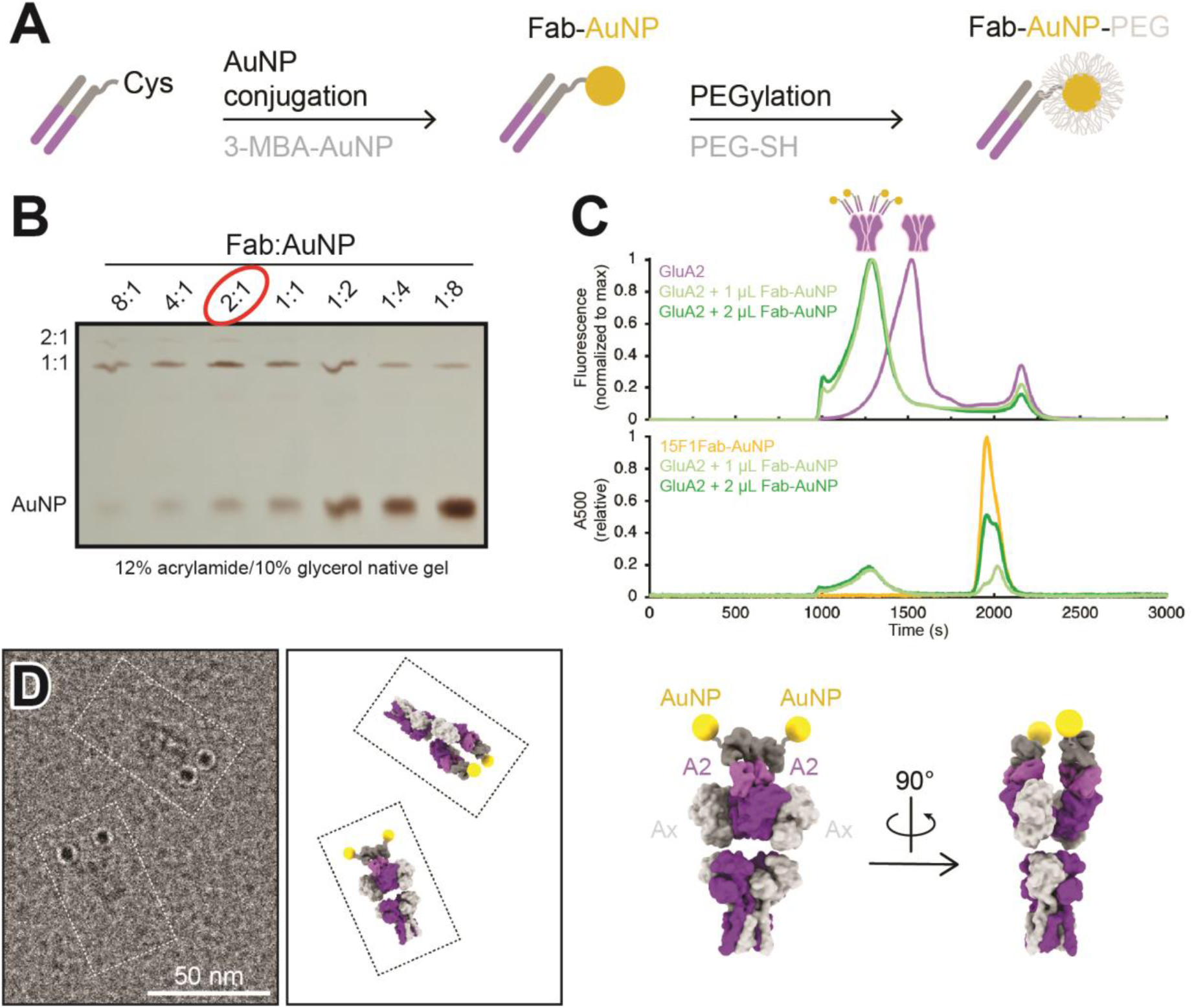
Development and characterization of anti-GluA2 Fab conjugated to AuNP. **(A)** Conjugation strategy for covalent linkage of AuNP to anti-GluA2 Fab. **(B)** Test-scale conjugations of Fab and AuNP at various Fab:AuNP ratios, run on native 12% acrylamide/10% glycerol gel. **(C)** Normalized FSEC traces of GFP-tagged GluA2 mixed with 1-2 µL of Fab-AuNP conjugate, measured at an excitation/emission of 480/510 nm (top; corresponding to GFP-tagged GluA2) or an absorbance at 500 nm (bottom; corresponding to AuNP). **(D)** Snapshot from single particle cryo-electron microscopy micrograph image showing two individual Fab-AuNP bound native mouse hippocampus AMPARs next to models depicting possible orientational views seen in image (PDB: 7LDD) (left). Model of anti-GluA2 Fab-AuNP bound to AMPAR with GluA2 in the B and D positions (right).

The anti-GluA2 15F1 Fab-AuNP conjugate remains fully capable of GluA2 binding, shifting a tetrameric GluA2 peak to an extent consistent with 1:1 Fab:GluA2 binding by fluorescence-detection size-exclusion chromatography (FSEC) [21] (Figure 1C). Taking advantage of an affinity tag included at the C-terminus of the Fab, the anti-GluA2 Fab-AuNP conjugate was used to affinity purify native GluA2-containing AMPARs from mouse hippocampal tissue [22, 23]. The isolated receptors were then visualized by single particle cryo-electron microscopy (cryo-EM) to investigate the stoichiometry of the 15F1 Fab-AuNPs bound to native AMPARs and to also determine the spatial positioning of the AuNPs relative to the receptor. Previous work from our group employing a similar anti-GluA2 15F1 Fab-directed purification of native hippocampal AMPARs revealed that the majority of GluA2-containing AMPARs in the mouse hippocampus contain two GluA2 subunits, occupying the B and D positions of the AMPAR complex [23]. In agreement with the studies of Yu and colleagues, imaging of the 15F1 Fab-AuNP-purified AMPARs revealed that the majority of particles harbor two clear AuNP densities present at positions consistent with the 15F1 Fab binding epitope on the amino terminal domain (ATD) of the GluA2 subunits (Figure 1D, Supplemental Figure 2). We do note that a minority of particles, less than about 20%, bear only a single AuNP, which may be due to either incomplete labeling, to labeling with a Fab devoid of an AuNP, to receptors with only a single GluA2 subunit, or to the unbinding of the Fab-AuNP and perhaps, separately, the loss of the AuNP itself. Nevertheless, we speculate that the majority of singly labeled receptors are simply those with one GluA2 subunit for several reasons. Not only was the Fab-AuNP conjugate purified by gel electrophoresis, but we also employed an excess of the Fab-AuNP conjugate, and the Fab-AuNP receptor complex was additionally purified by size exclusion chromatography.

### Fluorescence-guided localization of glutamatergic synapses

To enable the reliable, targeted imaging of AMPARs at excitatory glutamatergic synapses in the context of the relatively large area of the lamella, a transgenic mouse line expressing mScarlet fused to the presynaptic vesicular glutamate transporter 1 (vGlut1) and EGFP fused to the postsynaptic density protein 95 (PSD95) was generated (Figure 2A). The vGlut1-mScarlet mouse line was designed in an identical manner to a previously established vGlut1-mVenus knock-in line in which the C-terminus of vGlut1 is fused to mVenus [24]. vGlut1 is abundantly expressed in glutamatergic synaptic vesicles, allowing for clear fluorescent visualization of excitatory presynaptic sites within tissue [24]. At postsynaptic sites, PSD95 is the most abundant scaffolding protein within the postsynaptic density [25, 26] and thus we generated a mouse line in which PSD95 is tagged at the C-terminus with EGFP. To enable simultaneous visualization of pre- and postsynaptic sites by fluorescent imaging, the homozygous vGlut1-mScarlet and PSD95-EGFP mouse lines were bred to make a double transgenic mouse line expressing both vGlut1-mScarlet and PSD95-EGFP. Assessment of vGlut1-mScarlet and PSD95-EGFP expression by FSEC confirmed the presence of a major peak corresponding to each fusion protein in solubilized mouse whole brain tissue (Figure 2B). Therefore, we expect stable expression and minimal cleavage of the fusion proteins within the vGlut1-mScarlet/PSD95-EGFP mouse line. We note that the vGlut1-mScarlet/PSD95-EGFP mouse line has no apparent phenotype, and all animals exhibit wild-type growth, breeding and overall behavior, thus leading us to conclude that their glutamatergic synapses are representative of those in wild-type animals.

**Figure 2.**
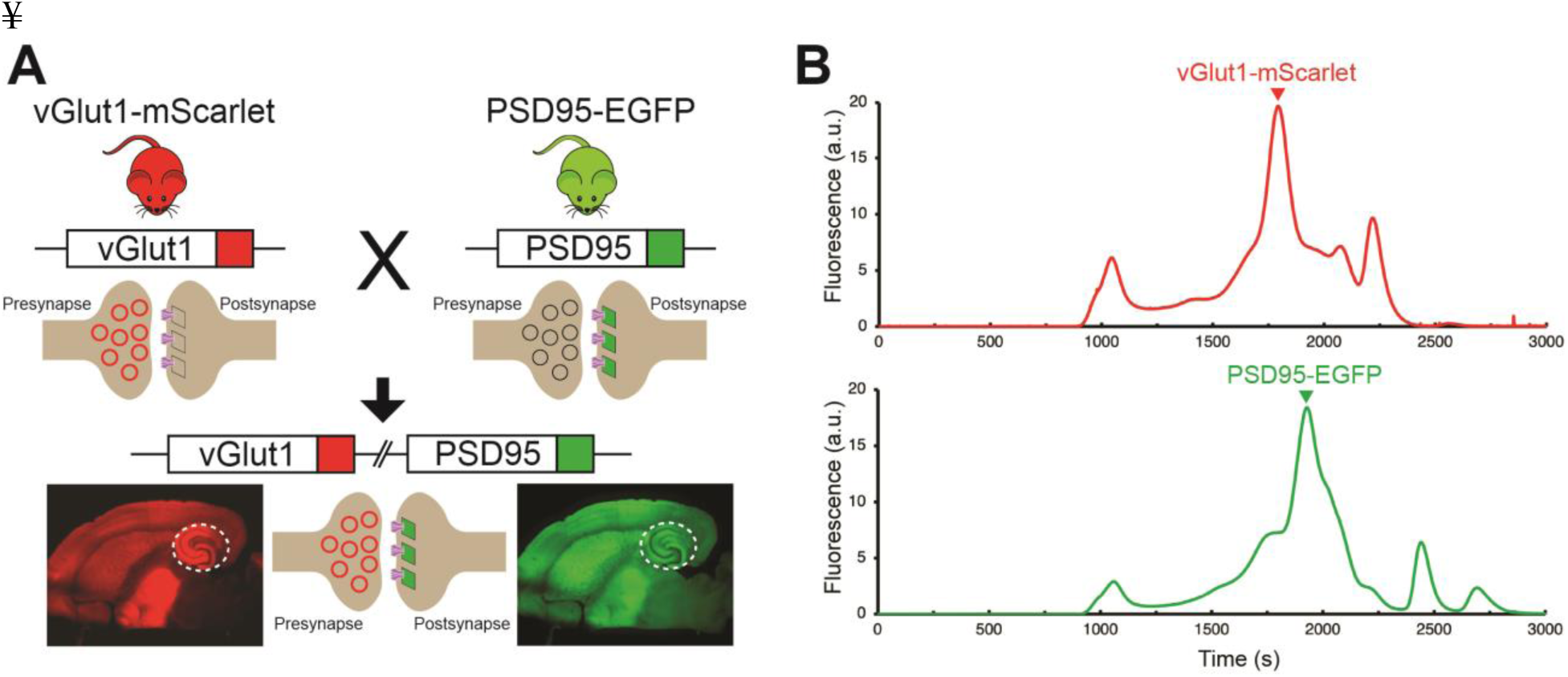
Development and characterization of transgenic mouse line for synaptic targeting. **(A)** Red fluorescent protein, mScarlet, was inserted at the C-terminus of vGlut1, which is present within synaptic vesicles at excitatory presynaptic terminals. In another mouse line, green fluorescent protein, EGFP, was inserted at the C-terminus of PSD95, which is highly expressed at excitatory postsynaptic sites. Homozygous vGlut1-mScarlet and PSD95-EGFP mice were crossed to generate a mouse line expressing both vGlut1-mScarlet and PSD95-EGFP to facilitate synapse identification. The white circle represents the hippocampus region. **(B)** Assessment of vGlut1-mScarlet and PSD95-EGFP in solubilized mouse whole brain tissue by FSEC confirms expression of each fluorescent fusion protein.

### Mouse hippocampus brain slice preparation and freezing

To visualize AMPARs at glutamatergic synapses in their native context, we developed a strategy to prepare and image mouse hippocampal brain tissue by cryo-ET. Ultra-thin horizontal brain slices from the double transgenic mouse line (vGlut1-mScarlet/PSD95-EGFP) were prepared using a vibratome set to 40 µm thickness (Figure 3A). Brain slices of this size are thin enough to minimize ice formation and make cryo-focused ion beam (cryo-FIB) milling less challenging, while thick enough to handle during immunolabeling and high pressure freezing. From each horizontal brain slice, the hippocampal region was excised carefully with a scalpel knife (Figure 3B). These hippocampal slices were incubated with the anti-GluA2 15F1 Fab-AuNP conjugate for 1 hour at room temperature on an orbital shaker. Following washout of the antibody-containing buffer, hippocampal slices were trimmed to isolate the CA1 region, which contains the Schaffer collateral pathway from CA3 synapses onto CA1 apical dendrites in the stratum radiatum (dashed line in Figure 3B) [27]. The slices were then briefly incubated in a buffer supplemented with 20% dextran, as a cryoprotectant. Two hippocampal CA1 slices were placed on the grid bar side of an electron microscopy (EM) grid coated with a 20-30 nm thick carbon film, with apical dendrites from CA1 neurons facing toward the center of the grid (Figure 3C). We note that the stratum radiatum shows brighter fluorescence both in red and green channels compared to the striatum lacunosum-moleculare (Figure 3B insets), which helped guide targeting.

**Figure 3.**
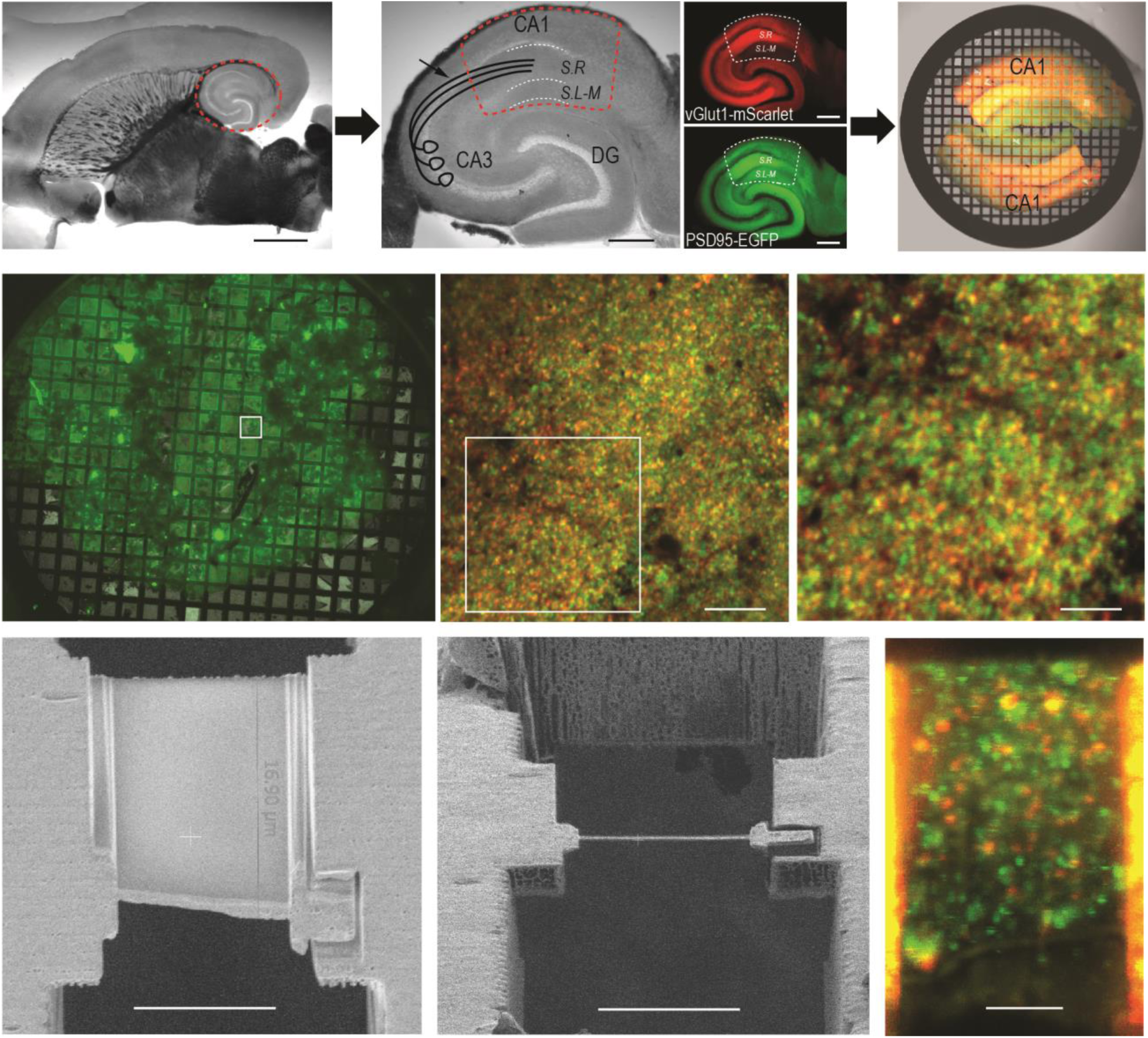
Hippocampus brain slice preparation and FIB milling of lamella. **(A)** Ultra-thin brain slice made from a vGlut1-mScarlet and PSD95-EGFP mouse. The red dashed circle indicates the hippocampus. Scale bar: 2 mm **(B)** A wide field image of excised hippocampus brain slice. The image also illustrates CA3 axons projecting to CA1 apical dendrites in the Schaffer collateral (arrow). Fluorescent images show vGlut1-mScarlet (top) and PSD95-EGFP (bottom) from the same hippocampus slice on the left. The dashed line (red or white) area indicates the region of interest for FIB milling (CA1 region of hippocampus), which is further trimmed with a scalpel knife. S.R: Stratum Radiatum, and S.L-M: Stratum lacunosum-moleculare. Scale bar: 500 µm **(C)** Overlay image of two trimmed hippocampus brain slices on EM grid, before high pressure freezing. The white dashed line indicates the stratum radiatum, the area with brightest fluorescence in the CA1 region and where the Schaffer collateral projects. Scale bar: 500 µm **(D)** A wide field green fluorescence image of a sample grid after high-pressure freezing. **(E)** Cryo-confocal image from a sample grid showing both vGlut1-mScarlet (red) and PSD95-EGFP (green) puncta. **(F)** Zoomed image of area indicated in E. White arrows point to colocalization of red and green fluorescent signals and indicate potential synapses. **(G)** SEM image of polished lamella. **(H)** FIB image of same lamella in G. **(I)** Cryo-confocal image of lamella. White arrows at fluorescence colocalization points indicate examples of regions targeted in tomography data collection.

We utilized the ‘waffle’ method [18] for high pressure freezing. At the preparation stage of the high pressure freezer, a sample grid was placed between the flat sides of two polished and pretreated planchettes. Immediately before high pressure freezing, gold fiducials prepared in media containing dextran and sucrose cryoprotectants were applied to the sample on the planchettes. During high pressure freezing, the tissue was pressed in between the grid bars and frozen. Because the sample thickness is approximately equivalent to the height of the grid bars, the portion of the sample that is between the grid bars is minimally compressed during freezing. High pressure freezing using the methods described above limited the formation of crystalline ice and results in cryo-fixed tissue without the introduction of substantial visible freezing artifacts.

### Saturation of binding sites by the 15F1 Fab-AuNP conjugate

To examine the extent to which the anti-GluA2 Fab-AuNP conjugate saturates available binding sites within the brain tissue slices, hippocampus slices were soaked with or without the 15F1 Fab-AuNP conjugate at a concentration of 40 nM for 30 mins. After washout, the tissue was subsequently soaked with the 15F1 Fab labeled with Janelia Fluor 646 (40 nM) for another 30 mins. We reasoned that if the 15F1 Fab-AuNP conjugate saturated all available binding sites on GluA2-containing AMPARs, further labeling of GluA2 containing AMPARs with the 15F1 Fab-Janelia Fluor 646 conjugate should be low. In tissue samples without 15F1 Fab-AuNP pretreatment, we observe clear 15F1 Fab-Janelia Fluor 646 signal that slightly overlaps with the PSD95-EGFP and vGlut1-mScarlet fluorescent signals (Supplemental Figure 3). However, in slices previously soaked with 15F1 Fab-AuNP, the 15F1 Fab-Janelia Fluor 646 signal was substantially reduced. Based on this experiment and the established propensity of the 15F1 antibody and its fragments to label GluA2-containing AMPARs [14], we expect that the ‘staining’ with the 15F1 Fab-AuNP used in our tissue preparation pipeline labels the majority of accessible GluA2-containing AMPARs within the tissue.

### Cryo-FIB milling of mouse hippocampus tissue

High-pressure frozen mouse hippocampus tissue slices on EM grids were loaded into an Aquilos 2 cryo-FIB/SEM microscope and milled using the ‘waffle’ method [18]. Scanning electron microscopy (SEM) images were aligned with fluorescent light microscope images of the entire sample grid and regions with maximal green and red fluorescence were identified for targeting. In the apical dendrites of CA1 neurons, there are abundant excitatory synapses, especially from CA3 axons to CA1 dendrites. Before freezing, we intentionally oriented the apical dendrites of CA1 neurons toward the center of the grid. By carrying out the FIB milling near the center of the grid, we increase the likelihood of targeting excitatory synapses instead of soma or distal dendrites (Figure 3C-F).

Waffle milling was most successful on grids where the grid bars were clearly visible in the FIB/SEM images, indicating sufficiently thin ice, and on grid regions where the surface was relatively smooth, as judged by FIB/SEM imaging. For each lamella, initial trench cuts were made to mill through the entire depth of the sample at the front and back edges of a lamella target site. After making trench cuts for each lamella target, typically amounting to about 10 per grid, the grids were tilted to a 20° milling angle and milled with progressively lower FIB current to a final thickness of approximately 150-250 nm (Figure 3G-H).

### Identification of synapses within lamellae by cryo-confocal imaging

To confirm the presence of intact synapses within lamellae and to guide synaptic targeting during cryo-ET data collection, lamellae were imaged by cryo-fluorescence confocal microscopy. Grids with milled lamellae were loaded onto a confocal microscope equipped with a cryo-correlative microscopy stage, thus maintaining the sample at cryogenic temperatures during imaging. Clear signals for both presynaptic vGlut1-mScarlet and postsynaptic PSD95-EGFP were visible on the majority of lamellae. Ideal synaptic targets on lamellae are indicated by colocalization of red presynaptic vGlut1-mScarlet and green postsynaptic PSD95-EGFP puncta, visualized as yellow overlapping spots (Figure 3E-F). At each lamella site, confocal fluorescence images were taken from the lamella area, typically about 20 x 13 µm in size. Because the lamellae were milled at a 20 degree angle, approximately eight 1 µm increment Z-stack images were taken and compiled to visualize an entire single lamella (Figure 3I). On an average single lamella, we typically detect at least a dozen spots with colocalized fluorescent puncta, highlighting ideal regions for synaptic targeting during subsequent cryo-ET data collection (Figure 3I, Supplemental Figure 4).

### Cryo-electron tomography of hippocampal synapses

Cryo-FIB milled lamellae were imaged using a 300 keV Krios cryo-transmission electron microscope. Initially, medium magnification montage maps of each lamella were taken to guide picking of synaptic targets. Often, clear cellular compartments containing synaptic vesicles can be identified by eye from the medium magnification EM maps. Additionally, cryo-correlative light and electron microscopy (cryo-CLEM) further guided the selection of targets (Supplemental Figure 4). Registration of the fluorescence and electron microscopy maps was performed by alignment of manually identified lamella structural features, often based on the edge shape of the lamella visible in both the confocal reflection and EM images. Once aligned, the presynaptic vGlut1-mScarlet signal often overlaps with cellular compartments containing synaptic vesicles in the EM map, substantiating alignment of the light and EM images. Synaptic targets were picked both by manually scanning the EM map for presynaptic vesicle-looking features as well as the overlaying fluorescence signal. Typically, we selected synaptic targets oriented such that the pre- and postsynaptic terminals are side by side with a visible synaptic cleft in between (Figure 4A-B). We picked as many as 10-30 tomography targets within a single 20 µm x 13 µm lamella depending upon the presence of synaptic features and quality of lamella ice. The tilt series of the targets were acquired using a grouped dose-symmetric scheme [28] starting normal to the lamella surface at - 20° and ranging from -68° to 28° with a 3° increment. The magnification was set to give a pixel size around 2 Å and data were collected at a defocus of -2.5 µm. For images collected normal to the lamella surface, the contrast transfer function (CTF) could be fit up to 5.6 to 7.0 Å (Supplemental Figure 5). Tomograms were reconstructed from tilt series taken at synaptic targets by patch tracking methods (Figure 4C). Within reconstructed tomograms, we observed the anticipated histological features of neuronal tissue, including myelin-wrapped compartments, microtubules, synaptic vesicles, and mitochondria (Figure 4D) [29]. Moreover, there were clusters of AuNPs nearby presynaptic vesicle-containing compartments, and a scattering of more dispersed AuNPs adjacent to the dense clusters. These AuNP features facilitated the identification of synaptic clefts, characterized by a 20-30 nm distance separating the pre- and postsynaptic membranes (Figure 4C).

**Figure 4.**
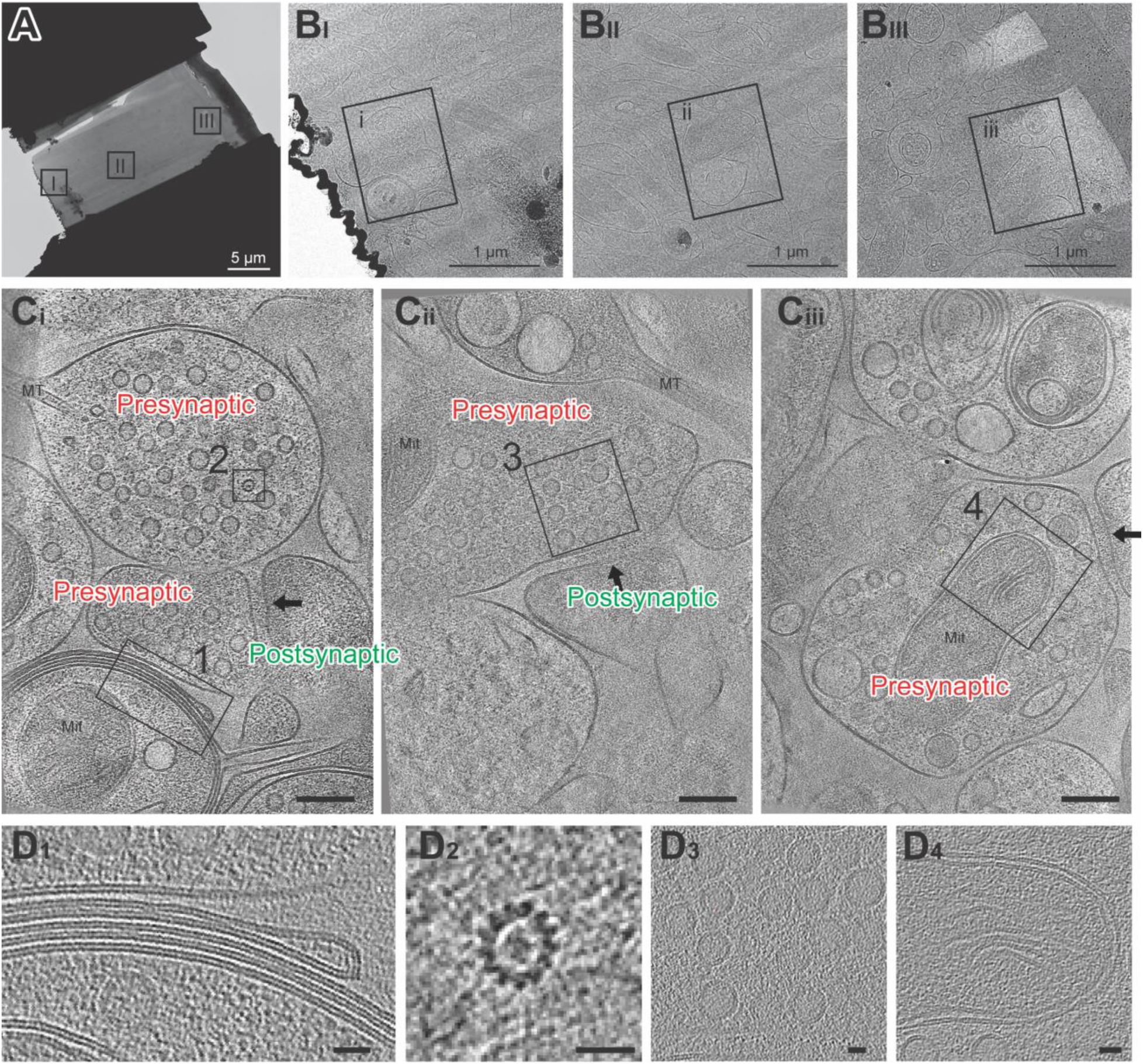
Cryo-electron tomography imaging of glutamatergic synapses within brain tissue. **(A)** Medium montage map image of lamella. **(B _I-III_)** Enlarged images of lamella indicated in A. **(C _i-iii_)** Slices from SIRT filtered tomograms collected at squares indicated in B. The synaptic cleft in each tomogram is indicated with a black arrow. Mit: mitochondria and MT: microtubule side views. One gold fiducial is visible in C _iii_. Scale bars are 100 nm. Each slice shown is about 10 nm thick. **(D)** Example image of myelin membranes (D_1_), top down view of microtubule (D_2_), synaptic vesicles (D_3_), and part of a mitochondria (D_4_). Scale bars are 20 nm. Each slice shown is 10 nm thick.

### Anti-GluA2 Fab-AuNP defines AMPARs at glutamatergic synapses

To better understand the spatial distributions of AuNPs bound to a single AMPAR in our tomograms, we first analyzed the positions of AuNPs in a single particle dataset of anti-GluA2 15F1 Fab-AuNP bound to native mouse hippocampus AMPARs. By preparing cryo-EM grids at a low particle density that clearly allowed for separation of individual receptor complexes, we ensured that nearest neighbor AuNP positions were attributable to AuNPs within, rather than between, receptor complexes. In turn, this allowed us to measure the nearest neighbor projection distance observed between AuNPs bound to an individual AMPAR (Figure 5A, Supplemental Figure 2). Throughout sample preparation we included the AMPAR antagonist ZK-200775 [30] and the positive modulator RR2b [31] to stabilize AMPARs in a resting, non-desensitized state [32]. The distance between the last structured residue (K216) in the heavy chain of each of the 15F1 Fabs in previously published structures of 15F1 Fab-bound hippocampal AMPAR in the resting state is approximately 80 Å (Supplemental Figure 6). In the 15F1 Fab-AuNP conjugate, the AuNP is covalently bound to a cysteine on the heavy chain located 11 residues after K216, allowing the AuNP as much as 35 Å of displacement from this point. As a result, inter-AuNP distances from Fabs bound to a single receptor can conceivably range from 30-150 Å, where an inter-AuNP distance of 30 Å would correspond to two ∼30 Å diameter AuNPs immediately adjacent to one another.

**Figure 5.**
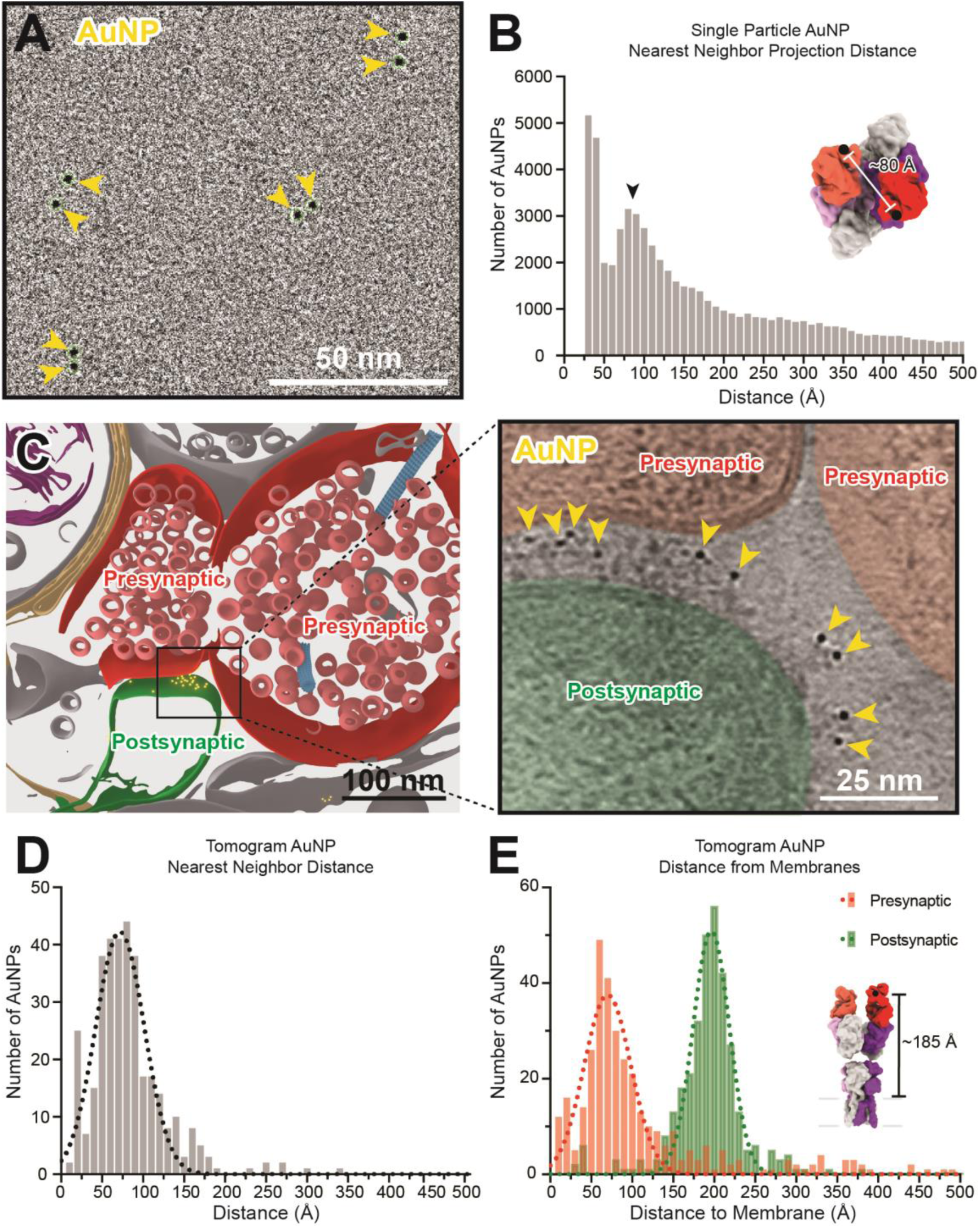
Anti-GluA2 Fab-AuNP labeling defines AMPAR position in single particle cryo-electron and cryo-electron tomography data. **(A)** Snapshot from low defocus single particle cryo-electron microscopy dataset showing four pairs of Fab-AuNPs bound to native mouse hippocampus AMPAR. **(B)** Quantitation of nearest neighbor AuNP projection distances from full ∼1000 micrograph dataset from (A), overlayed with Gaussian fit of data. Distance between structured C-termini of Fabs in hippocampal AMPAR structure (PDB: 7LDD) shown for reference. **(C)** Segmentation (left) and slice through tomogram (right; denoised using IsoNet) of mouse hippocampus CA1 tissue stained with anti-GluA2 Fab-AuNP, with AuNPs visible in-between the pre- and postsynaptic membranes. Quantitation and Gaussian fits of nearest neighbor AuNP distances **(D)** and distance to the closest point on the pre- or postsynaptic membrane **(E)** from five Fab-AuNP labeled synapses. Distance between structured C-terminus of Fab and the putative membrane location in hippocampal AMPAR structure (PDB: 7LDD) shown for reference.

The majority of nearest neighbor AuNP projection distances measured in the single particle dataset are in agreement with this range, with major peaks present around 40 and 85 Å (Figure 5B). This analysis is based on projection distances, which will generally underestimate the actual distance between AuNPs and will undercount AuNPs overlapping in Z with an inter-AuNP projection distance of less than one AuNP diameter (which are either measured at the minimum picking distance of 30 Å apart or detected as a single AuNP). Additionally, this analysis is subject to the effects of preferred orientation bias and aggregation of AMPARs on single particle cryo-EM grids. Despite these challenges, the histogram of observed inter-AuNP distances has a major peak at 85 Å, which is in overall agreement with the expected 2:1 Fab-AuNP:AMPAR stoichiometry and geometry. These analyses provided confidence that our anti-GluA2 Fab-AuNP conjugate labels hippocampal AMPARs in their native state without overall disruptions to AMPAR structure.

Careful inspection of the synaptic cleft in our tomograms revealed a plethora of strong ∼30 Å diameter densities (Figure 5C), consistent with the presence of 15F1-AuNP label. We then used automated tools to create segmentations for the positions of AuNPs and membranes as described in the methods. Nearest neighbor distances of AuNPs at synapses fall mostly within the range of 40-100 Å, fitting to a Gaussian mean of 72 ± 30 Å (Figure 5D). AuNPs within tomograms are often densely clustered at putative AMPAR nanodomains [33], likely contributing to a slight reduction in observed nearest neighbor distances in comparison to the single particle data, in which the low density of AMPARs maximizes analysis of AuNPs bound to individual rather than adjacent AMPARs. The distance between the 15F1 Fab heavy chain K216 residue and the putative membrane in previously published native AMPAR structures is about 185 Å, where the extracellular boundary of the membrane was estimated at a plane defined by four pre-M1 helix residues (Supplemental Figure 6). Within 15F1 Fab-AuNP-labeled tomograms, synaptic AuNPs are located at a mean distance of 196 ± 22 Å from the postsynaptic membrane and 70 ± 29 Å from the presynaptic membrane (Figure 5E), clearly defining the position of AMPARs at synapses (Supplemental Video 1).

## Discussion

The overall goals of the present work are to develop methods to enable the identification and localization of the molecular machinery at chemical synapses in unstained, unfixed native brain tissue slices in order to place mechanisms of synaptic transmission, plasticity and development on a 3D structural foundation. By harnessing genetically engineered mouse lines, together with cutting of thin brain slices and their subsequent HPF and FIB milling, we have produced substrates for cryo-ET that allow us to target glutamatergic synapses by cryo-fluorescence microscopy at micrometer resolution. Because we have, in parallel, labeled the predominate GluA2 subunit of AMPARs with a highly specific Fab-AuNP conjugate, we can identify glutamatergic synapses and AMPARs at nanometer resolution. This labeling enables clearer identification of synapses and receptor positioning compared to prior studies using cultured neurons or synaptosomes [9, 11, 12]. These methods will not only facilitate the specific visualization of the molecular machinery of excitatory synapses, but also because they are generalizable, we imagine that similar strategies will be useful for identifying and localizing other key molecules at glutamatergic synapses, such as *N*-methyl-D-aspartate receptors, calcium channels and vesicle fusion machinery. The approaches outlined here may also be helpful in revealing the macromolecular organization of a wide range of different synapses and cellular structures.

Despite the advances enabled by the approaches described here, there are nevertheless several limitations of the present work. Firstly, while we endeavored to target CA3-CA1 synapses, our ability to accurately and reproducibly identify pyramidal cell-pyramidal cell synapses remains limited, in part because of the resolution limitations of the cryo-fluorescence microscopy and the much smaller field of view in the transmission electron microscopy. Further efforts will be required to optimize correlative targeting and faithful identification of specific synapses, which may include development and use of additional fluorescence or electron dense labels. We also are aiming to better understand the extent to which the labeling of the GluA2-containing receptors is complete and if not, how to ‘push’ the labeling to completion. The extent to which labeling of GluA2-containing receptors with the Fab AuNP conjugate alters the interactions of the receptors with synaptic proteins, and thus synaptic architecture and function, also warrants further investigation. In addition, the step toward subtomogram averaging of GluA2-containing AMPA receptors, using the AuNP labels, may ultimately be hindered by the flexible linker between the Fab and the AuNP, thus requiring the development of new reagents in which the AuNP is rigidly bound to the Fab, or subtomogram averaging approaches that do not rely upon the AuNP positions. Lastly, a crucial process in the preparation of brain slices for HPF presently involves inclusion of cryoprotectants in the media, in order to minimize ice formation. At the present time we do not know how the cryoprotectants alter the functional state of the tissue and the synapses. Further experiments will be required to determine if we can utilize alternative cryoprotectants or reduce their use. If they remain essential, then we aim to better understand their effects on the functional states of the synapses.

Upon resolving limitations of the methods described here, and working further toward optimization of sample preparation, cryoprotection, targeting and milling, in combination with ideal collection of tilt series and image processing, we hope to carry out subtomogram averaging of synaptic receptor complexes. We further aim to identify additional components of the synaptic machinery and pursue their reconstructions, ultimately positioning ourselves to compare the structures of synapses from different brain regions, at selected times during development or in states of disease, thus furthering our understanding of the interrelationships between synapse structure and function.

## Acknowledgements

We thank Dr. Lauren Ann Metskas (LAM; Purdue) for advice, education and training related to cryo-electron tomography methods, Dr. Chang Sun for assistance with software installation and computer workstations, Dr. Nelson Spruston (HHMI Janelia) and the HHMI Janelia transgenic facility for their support in generating the precursor line of the PSD95-CreNABLED2 mouse, Rachel Courtney for assistance with manuscript preparation, Mark Mayer and LAM for comments on the manuscript, and Dr. John T. Williams for sharing a vibratome. We acknowledge the generous support and use of the HHMI Janelia Cryo-EM facility for FIB milling on an Aquilos2 and data collection on Krios microscopes, which were operated by Drs. Shixin Yang, Rui Yan, and James Jung (HHMI Janelia), and the OHSU MMC facility for data collection on the Glacios with assistance from Erin Stempinski. C.J.S. is supported by grant number CA253730 from the National Cancer Institute at the National Institutes of Health. E.G. is an investigator of the Howard Hughes Medical Institute and gratefully acknowledges the generous support of Jennifer and Bernard LaCroute.

## Author contributions

A.M. and C.J.S. prepared samples and performed the experiments. A.M, C.J.S, and J.E. processed and visualized data. A.M. and C.J.S., together with E.G., designed the project and wrote the manuscript. M.S. and N.J. performed cryo-FIB milling of samples. X.Z. and Z.Y. assisted with tilt series data acquisition. M.Q. and H.Z. provided the PSD95-EGFP mouse line.

## Data availability

Tomograms have been deposited to EMDB under accession codes EMD-44174, EMD-44175, and EMD-44176. The corresponding raw data has been deposited to EMPIAR under accession code EMPIAR-11984.

