## Supplemental Material for "Cryo-electron tomographic investigation of native hippocampal glutamatergic synapses"

### Methods

#### Dissection and preparation of hippocampal slices

All procedures were performed in accordance with the guidelines of Oregon Health & Science University and the Animal Care and Use Committees approved all of the experimental procedures.

We prepared a mouse line in which we appended mScarlet [34], a red fluorescent protein, at the C-terminus of vGlut1, based on a previously created vGlut1-mVenus mouse line. A second mouse line, where the C-terminus of PSD95 is tagged with EGFP, was a generous gift from Dr. Haining Zhong [25]. The mice used in the present study are a cross between the vGlut1-mScarlet (+/+) and PSD95-EGFP (+/+) lines and were subsequently bred to generate a vGlut1-mScarlet (+/-)/PSD95-EGFP (+/-) mouse line. Adult male and female mice (8-12 weeks old) were anesthetized with isoflurane and decapitated. Brains were quickly removed and placed in a vibratome (Leica, VT1200). Horizontal brain slices (40 µm) were prepared at room temperature in 95%$O_2$/5%$CO_2$ oxygenated HEPES base solution containing the following (in mM) 150 NaCl, 2.5 KCl, 2 $CaCl_2$, and 10 HEPES (pH 7.3). AMPA receptor antagonist, ZK-200775 (1 µM) and the positive allosteric modulator, RR2b (1 µM), were included throughout the experiments. From a horizontal brain slice, hippocampal regions were excised and incubated with the GluA2 subunit-specific 15F1 Fab conjugated with AuNP (2 µg/mL or ~40 nM) for 1 hour at room temperature on an orbital shaker (180 rpm), followed by 3 washes for 15 mins each with HEPES base solution at room temperature. Hippocampal slices were further dissected with a scalpel knife to isolate the CA1 region, which was then incubated in a HEPES base solution containing 20% dextran for at least 30 min prior to high-pressure freezing (HPF).

**HPF of hippocampal brain slices**

*Freezing preparation:* Planchettes and grids were pretreated such that frozen sample grids could be easily detached from the planchettes after freezing [18, 35]. The planchettes (Cu/Au 6 mm,Type B 0.3 mm cavity) were polished on the flat side, using 7000 and 15000 grit sandpaper, until the surface appeared shiny. The surfaces were then cleaned with a metal polish, Wenol. The flat sides of the planchettes were coated with 1-hexadecene for at least 30 min, which was removed right before use.  200 µL of PEGylated 10 nm gold fiducials (CP11-10-PA-1K-DI water-50, Nanoparts) were spun down at 15,000 rpm for 30 mins and the supernatant was removed. After adding 200 µL of HEPES base solution, the centrifugation step was repeated and the supernatant was again removed. Next, 50 µL of HEPES base solution containing cryoprotectants (20% dextran and 5% sucrose) was added. This mixture was stored at room temperature until use. Right before freezing samples, 200 mesh gold extra thick carbon grids (CF200-Au-ET; EMS) were glow discharged with a PELCO easiGlow unit (Ted Pella) for 30 s at 15 mA current with the grid bar sides up.

*HPF of brain slice sample:* The sample and 6 mm flat specimen planchettes were prepared for HPF at the loading station of a Leica EM ICE. First, a bottom planchette was placed on a lower half cylinder. Two CA1 tissue slices were placed on the grid bar side of a grid with apical dendrites facing toward the center of the grid, and a sample grid was placed on the bottom planchette sample side up. Right before HPF, a small volume (1.5-3 µL) of 10 nm diameter PEGylated gold fiducials (Nanoparts) in HEPES base solution supplemented with 20% dextran and 5% sucrose was applied on top of the sample grid. A second planchette was then placed on top of the sample, flat side down, making sure to not introduce an air bubble. The assembled cartridge was then subjected to the automated HPF cycle that involves pressurization to approximately 2050 bar (210 kPa) and cooling to -196˚C within 10 ms. After HPF, samples were transferred into the autogrid clipping station to disassemble the grid from the planchettes. Sample grids were then clipped using a cryo-FIB autogrid ring and C-clip, with the carbon layer facing the autogrid ring and the sample side facing the C-clip.

**Cryogenic focused ion beam (Cryo-FIB) milling**

The HPF hippocampal brain slice grids were milled on an Aquilos 2 cryo-FIB/SEM microscope (Thermo Fisher Scientific). Scanning electron microscope (SEM) images of the whole grid were taken using a nominal magnification of 350x and an acceleration voltage of 2 kV with a beam current of 13 pA and a dwell time of 1 µs. Sputter coating with a platinum layer (30 mA current for 15 sec at 10 Pa pressure) and a gas injection system (GIS) deposition of a trimethyl(methylcyclopentadienyl) platinum layer (2 min) were applied before FIB milling. The brain slices were placed on the grid such that the Schaffer collaterals, where CA3 axons synapse onto CA1 apical dendrites, were located near the center of the grids. We picked lamella positions in the middle of the grids to increase the likelihood of imaging these CA3 – CA1 synapses. The HPF brain tissue lamellae were milled using the 'waffle' method [18, 35]. Briefly, two rectangular trench cuts were milled at 15 nA ion beam current to clear the front and back of the target lamella site. After making trench cuts at each site, a preemptive "preclean" step was utilized to further remove the bottom portion of the sample. The precleans were completed by using incremental angles; starting at 40˚, then 30˚, followed by 20˚. Following precleans, the notch patterns were milled at the side of the final lamella sites. Automated milling was performed at the milling angle of 20˚ by progressively reducing the ion beam current from 1 nA to 30 pA to produce a final lamella thickness of approximately 150-250 nm. After automated milling, the grids were tilted and manually polished at both +0.5˚ (20.5˚) and -0.2˚(19.8˚) with an ion beam current of 50 or 30 pA. The milled grids were retrieved from the Aquilos 2 and stored in liquid nitrogen until they were imaged.

**Cryo-confocal imaging of milled sample grids**

The milled grids were loaded onto a confocal microscope (Zeiss LSM980) fitted with a CMS196 cryo-stage (Linkam). The chamber and the grid stages were maintained below -185˚C throughout imaging. First, the whole grid was imaged via widefield fluorescent light using 5x and 10x objectives. Fluorescence excitation was elicited using an X-Cite Xylis LED light (Excelitas technologies) and emission was detected using an Axiocam 506 camera (Zeiss). For higher resolution, confocal laser scanning microscope images were taken using a 100x (0.75 NA) objective with and without an Airyscan module. Imaging settings, such as pixel resolution and pinhole diameter, were set according to the Zen 3.7 software for optimal Nyquist-based resolution. Laser power settings and dwell times were set slightly higher and longer than those used for room temperature experiments, as the photobleaching is reduced at cryogenic temperature [36]. At this sample thickness, the laser exposure did not cause significant heating and thus the sample was not devitrified. At each lamella, a cropped imaging area was designated to include only the lamella area and a Z-stack range was set from the milling edge to the end of the lamella with a 1 μm interval Z-spacing. Approximately eight Z-stack images were captured and compiled into a maximum intensity projection to visualize an entire single lamella at -20° tilt.

**TEM tilt series acquisition**

Tilt series were acquired using a 300 keV FEI Titan Krios cryo transmission electron microscope, equipped a spherical aberration corrector, a Gatan BioContinuum energy filter (6eV energy slit width) and a K3 direct detector (Gatan, CA). Medium montage map images from each lamella were collected with a nominal magnification of 2,250x with a calibrated pixel size of 60.3 Å. Tilt series were collected using a dose-symmetric scheme [28] starting at -20° and ranging from -68° to 28°, with a 3° increment using SerialEM [37]. Data were acquired with a magnification of 33,000 (calibrated pixel size of 2.06Å) on a K3 detector with a total electron dose of 150 $e^-/Å^2$ and a target defocus of -2.5 µm. At each tilt, a movie of 8 frames was collected.

**Tomogram reconstruction with patch tracking**

Movies were motion corrected using MotionCor2 [38] without dose weighting and binned to a pixel size of 2.5 Å. Initially, tomograms were reconstructed using the fiducial-free alignment software AreTomo [39]. The initial tomograms were inspected individually to determine which tomograms warranted further image processing. Tilt series with low quality ice, large ice contamination, or with no discernable Fab AuNP particles, were discarded. Several good quality tilt series were chosen to go through tomogram reconstruction in Etomo using patch tracking [40]. Tilt series were binned by 4 for patch tracking alignment and CTF corrected with Ctfplotter within Etomo [41]. The final tomograms were binned to a final pixel size of 10 Å.

**AuNP synthesis**

Thiolate-protected gold nanoparticles (AuNPs) were prepared as previously described [15, 20] using a 7:1 molar ratio of 3-mercaptobenzoic acid (3-MBA):$HAuCl_4$ to produce uniform nanoparticles with diameters of approximately 3 nm. Briefly, 1 mL of 84 mM 3-MBA was mixed with 0.4 mL of 28 mM $HAuCl_4$ in a 15 mL plastic (Falcon) tube, and 3.5 mL of water was added, producing a white precipitate. NaOH was added dropwise to 100 mM (49.5 µL of 10 M NaOH), resolubilizing the precipitate. The reaction was mixed by rotation at room temperature for at least 16 hours, after which the solution was moved to a 50 mL plastic tube and 28.7 mL of 27% methanol was added. $NaBH_4$ was added to a final concentration of 2 mM, using approximately 450 µL of a 150 mM stock solution of $NaBH_4$, and the reaction was continued by rotation at room temperature for 4.5 hours. AuNPs were precipitated by adding NaCl to 100 mM (694 µL of 5 M NaCl added to a 34 mL reaction), followed by the addition of 80 mL of methanol. The reaction solution was distributed into three 50 mL plastic tubes and centrifuged at 4100 *g* for 20 minutes. Pellets were gently washed with 75% aqueous methanol, combined into a single plastic tube, and centrifuged at 4100 *g* for 20 minutes. The remaining methanol was removed and the pellet was dried overnight in a desiccator. The dried pellet was resuspended in water to ~10 mg/mL, resulting in a black homogeneous solution. An aliquot of AuNPs was diluted 1:1 with 10% aqueous glycerol in 0.2X TBE (20 mM Tris base, 20 mM boric acid, 0.5 mM EDTA) and run on a 10% glycerol, 12% PAGE gel in 0.2X TBE running buffer to assess AuNP size and homogeneity.

**Anti-GluA2 15F1 Fab-AuNP conjugation**

An anti-GluA2 15F1 Fab [14] construct was designed with an extended heavy chain fragment containing a single hinge cysteine for AuNP conjugation (C-terminal sequence:

KVDKKIVPRDAGAKPC) followed by a C-terminal Twin Strep tag. This construct, deemed 15F1FabxC-2xStr, was expressed in Sf9 insect cells by baculovirus transduction [23] and purified over Strep-Tactin Superflow resin in T150 buffer (20 mM Tris-Cl pH 8.0, 150 mM NaCl). Approximately 400 µg of 15F1FabxC-2xStr was diluted to 20 µM in TBE (100 mM Tris base, 100 mM boric acid, 2.5 mM EDTA) with 2 mM tris(2-carboxyethyl)phosphine (TCEP) and incubated at 37°C for 60 minutes. The reduced Fab was purified by size exclusion chromatography and concentrated to 1 mg/mL. To determine optimal AuNP conjugation conditions, a range of Fab:AuNP ratios were tested in small scale conjugation reactions. Fab (1 mg/mL; 1-8 µL) and AuNP (10 mg/mL; 1-8 µL) were mixed in TBE at a total volume of 10 µL (Fab:AuNP volumes of 8:1, 4:1, 2:1, 1:1, 1:2, 1:4, or 1:8 µL) and incubated at 37°C for 30 minutes before diluting 1:1 with 10% glycerol and loading 15 µL onto a 10% glycerol, 12% PAGE gel. A 2:1 Fab:AuNP ratio was determined to be the optimal condition for conjugation, and the reaction was scaled up by incubating 300 µL of 1 mg/mL 15F1FabxC-2xStr with 150 µL of 10 mg/mL AuNP and 50 µL 10X TBE at 37°C for 30 minutes. After adding 25 µL of glycerol, the entire reaction was run on a 10% glycerol, 12% PAGE gel (Supplemental Figure 1C) [15]. Bands corresponding to the 1:1 Fab:AuNP species were cut out, chopped into small pieces, and incubated overnight at 4°C in TBE buffer to extract the 15F1 Fab-AuNP conjugate. The supernatant containing the 15F1 Fab-AuNP conjugate was concentrated to 400 µL and incubated with 0.5 mM of PEG550-SH (3.3 µL of 60 mM PEG550-SH) for 60 minutes at 37°C. An optimal PEG550-SH concentration of 0.5 mM was determined to PEGylate the AuNPs without displacing the Fab (Supplemental Figure 1D). Excess PEG was removed by serial concentration and the final PEGylated Fab-AuNP was stored at 4°C. The final concentration of 15F1 Fab-AuNP was estimated by comparison to Fab standards on a denaturing gel.

**Hippocampal AMPAR purification with 15F1 Fab-AuNP**

Hippocampi from 10 male and 4 female adult vGlut1-mScarlet mice (total tissue mass = 675 mg) were resuspended in chilled homogenization buffer (20 mM Tris-HCl pH 8.0, 150 mM NaCl, 0.8 µM aprotinin, 2 µg/mL leupeptin, 2 µM pepstatin A, 2 µM ZK-200775, 2 µM JNJ-55511118, 50 µM RR2b) at 7.5 mL buffer per gram of tissue. Tissue was homogenized with 10 strokes on a Dounce homogenizer, then diluted 1:1 with solubilization buffer (homogenization buffer plus 4% w/v digitonin) and incubated at 4°C for 15 minutes on a nutator. The solubilized tissue was centrifuged at 4000 *g* for 3 minutes at 4°C and filtered through a 0.22 µm filter. Anti-GluA2 15F1 Fab-AuNP was added to the sample to a final concentration of approximately 40 nM and the sample was 'nutated' at 4°C for 15 minutes. The solution was then passed over 5 mL of StrepTactin Superflow resin equilibrated in purification buffer (20 mM Tris-HCl pH 8.0, 150 mM NaCl, 2 µM ZK-200775, 2 µM JNJ-55511118, 1 µM RR2b, 0.075% w/v digitonin) by gravity flow and eluted with purification buffer + 5 mM desthiobiotin. The resulting native AMPAR bound to 15F1 Fab-AuNP was further purified by size exclusion chromatography, using an HPLC. The absorbance at 280 nm was monitored and peak fractions corresponding to the 15F1 Fab-AuNP bound native AMPAR species were collected by hand and concentrated to 30 µL. The concentrated sample was diluted 1:1 in purification buffer and 3 µL were applied to glow discharged Quantifoil R2/1 Cu 200 mesh grids covered with a 2 nm continuous carbon film. The grids were blotted and plunge frozen by submersion into a 35/65% ethane/propane mix using an FEI Vitrobot set to 4°C, 100% humidity, 30 s wait time, 2.5 s blot time, and 0 blot force.

### Single particle cryo-EM data collection and processing

Single particle cryo-EM grids with native mouse hippocampal AMPAR bound to 15F1 Fab-AuNP were imaged on a 200 kV Thermo Fisher Glacios microscope equipped with a Gatan K3 Summit direct electron detector at a magnification of 45,000x (pixel size of 0.45 Å in super-resolution mode) and a total dose of 50 e/$Å^2$. Data were collected using the SerialEM multi-shot 3x3 pattern with each movie containing 50 frames collected over a total exposure time of 2.9 s. For analysis of nearest neighbor inter-AuNP distances, a "low defocus" dataset of approximately 1,000 movies was collected at a defocus of -1.0 to -1.2 µm, a range in which the automated AuNP bead picking software 'imodfindbeads' [40] was most accurate at picking AuNPs and avoiding background signal, as judged by manual analysis of picking results. For optimal visualization of receptor density along with AuNPs, several movies were collected at a higher defocus range of -4.5 to -5.0 µm. All movies were motion corrected in cryoSPARC with patch motion correction [42].

### Tomogram segmentation and AuNP distance analyses

A custom python script was written to identify the 2D coordinate positions of AuNPs from all motion corrected single particle cryo-EM movies in the "low defocus" dataset and calculate the nearest neighbor distance to the next closest AuNP for each AuNP position. The 'imodfindbeads' command within IMOD (version 4.12.56) was used for automated identification of AuNPs within the micrographs [40].

To analyze inter-AuNP and AuNP-membrane distances within tomograms, membranes in each tomogram were segmented with membrain-seg [43]. The 'findbeads3D' command within IMOD (version 4.12.56) was used to identify the 3D coordinate positions of all AuNPs from within a manually specified subregion of the tomogram, using the following options: BeadSize = 1.5, ThresholdForAveraging = 20, and StorageThreshold = 0.5. An optimal thresholding value to accurately pick AuNPs over background was manually determined for each analyzed subregion. The identity of presynaptic, postsynaptic, and myelin membranes were manually annotated and microtubules were traced by hand. The distances to the nearest AuNP neighbor and point on the pre- or postsynaptic membrane were calculated for each identified AuNP and fit to Gaussian distributions by nonlinear regression analysis using GraphPad Prism 10.1.1. Segmentations were rendered using Blender and the MolecularNodes plugin [44] . Where indicated, tomograms were denoised using CryoCare [45] or IsoNet [46].

**Supplemental Figures**

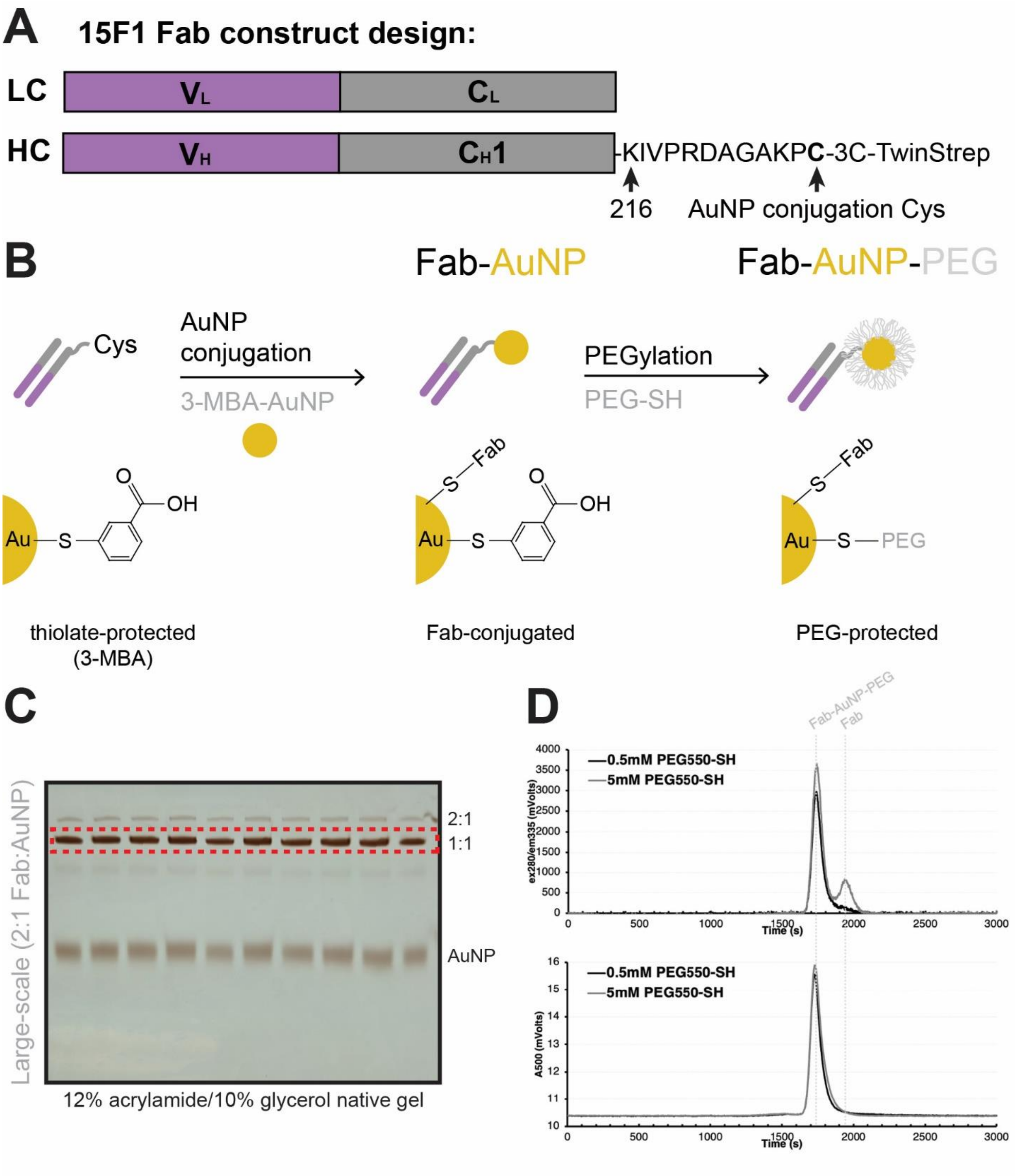

**Supplemental Figure 1. Preparation of PEGylated AuNP-15F1 Fab conjugate. (A)** Design of anti-GluA2 15F1 Fab construct for AuNP conjugation. The light chain (LC) is included without modification, while the heavy chain contains an extension of the constant domain 1 to include a single hinge cysteine for AuNP conjugation, followed by a 3C protease cleavage site and Twin-Strep tag. **(B)** Strategy for covalent conjugation and subsequent PEGylation of anti-GluA2 15F1 Fab and 3-MBA-protected AuNPs. **(C)** Full-scale conjugation of 15F1 Fab and AuNP using 2:1 Fab:AuNP ratio, with entire reaction run on native 12% acrylamide/10% glycerol gels (representative shown). The gel bands cut out for subsequent purification are denoted by the dashed red box. **(D)** Assessment of AuNP PEGylation with 0.5 or 5 mM PEG550-SH by SEC, using HPLC measurement of tryptophan fluorescence (top) and AuNP absorbance at 500 nm (bottom).

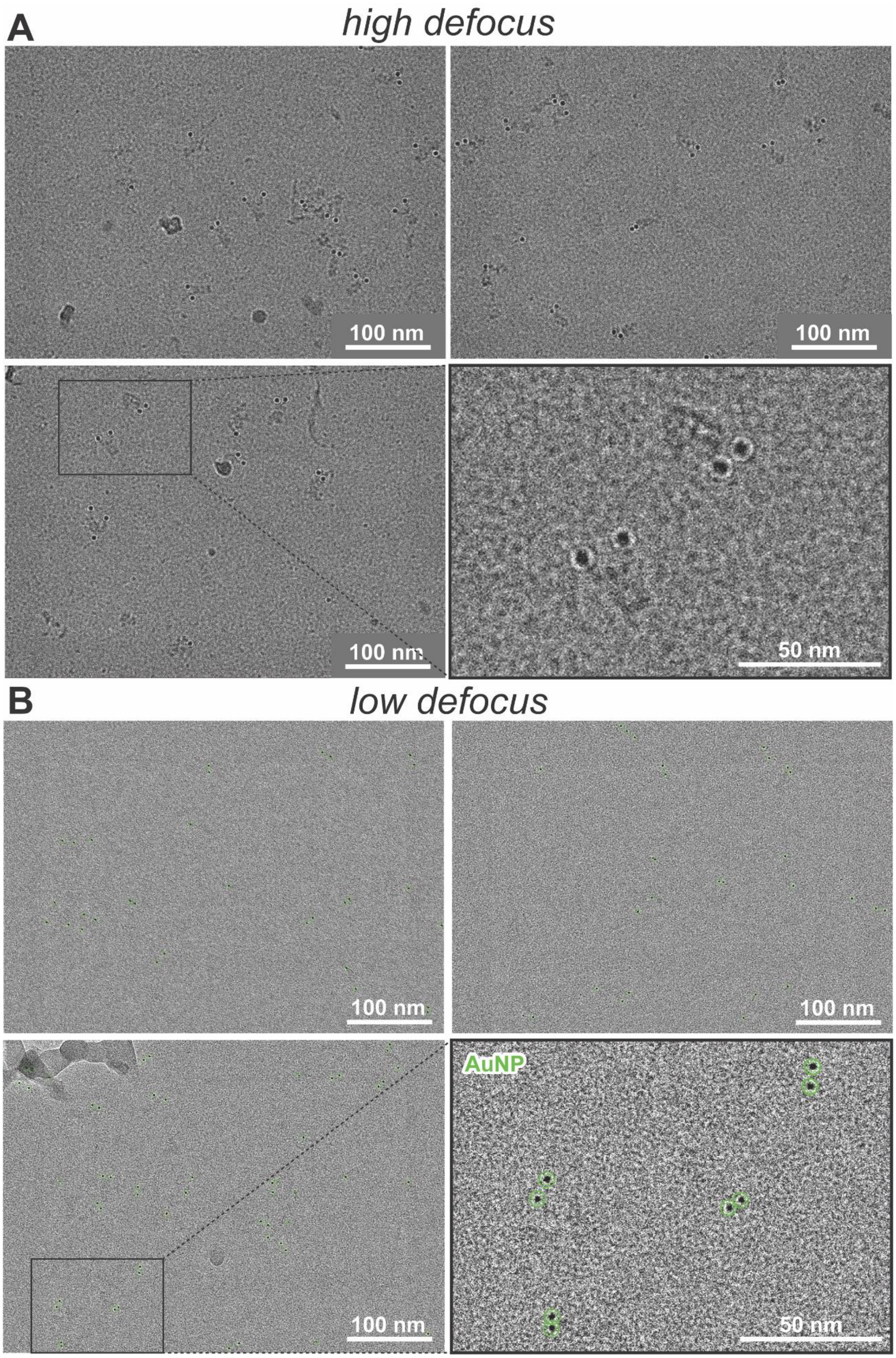
A
high defocus
100 nm
100 nm
100 nm
50 nm
B
low defocus
100 nm
100 nm
AuNP
100 nm
50 nm

**Supplemental Figure 2. Single particle cryo-EM of Fab-AuNP bound to native mouse hippocampal AMPAR.** Example single particle cryo-EM micrographs of native mouse hippocampal AMPAR bound to anti-GluA2 15F1 Fab-AuNP taken at **(A)** “higher” (-4.5 to -5.0 µm) and **(B)** “lower” (-1.0 to -1.2 µm) defocus ranges. Example automated bead picking hits from low defocus micrographs denoted with green circles.

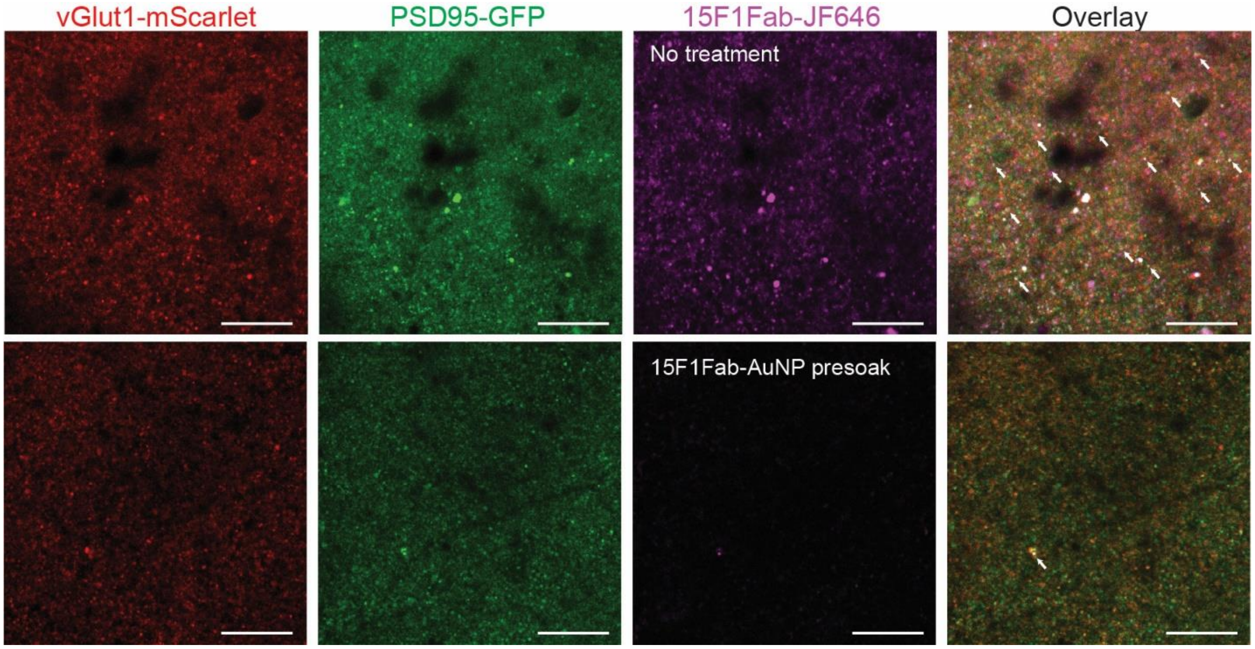


**Supplemental Figure 3. Pre-soaking tissue with 15F1 Fab-AuNP blocks majority of available GluA2 binding sites.** Blocked staining experiment in which mouse hippocampus tissue slices were either left in buffer (top) or presoaked with 15F1 Fab-AuNP (bottom) before subsequent staining with Janelia Fluor 646 (JF646)-labeled 15F1 Fab. Representative images from each sample are shown. Scale bars are 20 µm.

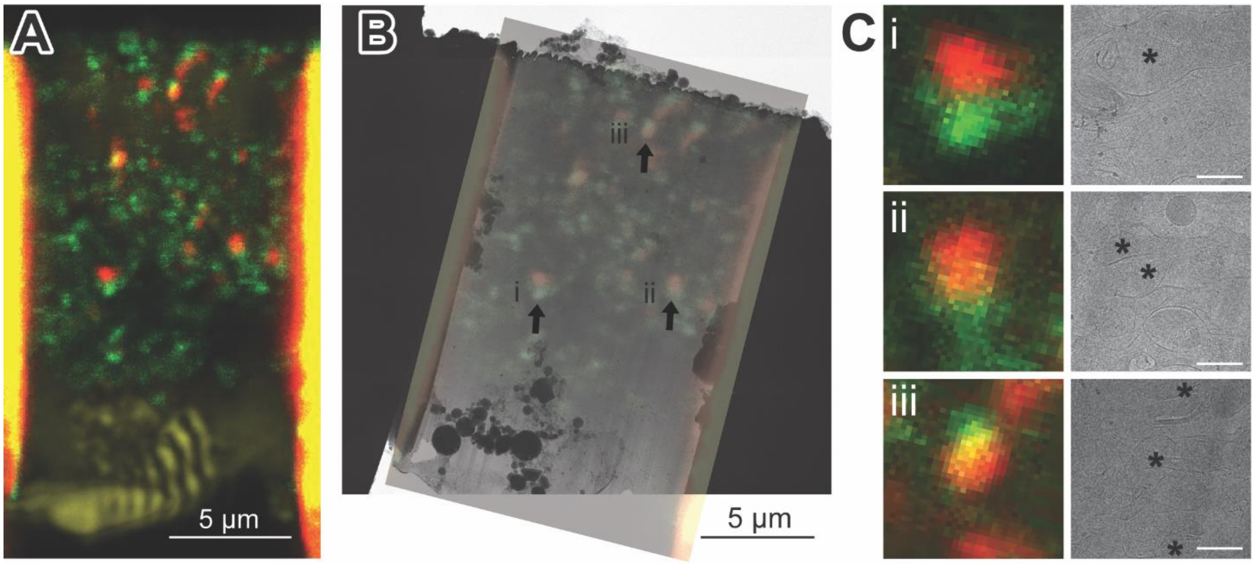


**Supplemental Figure 4. Cryo-CLEM guides synaptic targeting on lamella. (A)** Cryo-confocal image of lamella showing vGlut1-mScarlet (red) and PSD95-EGFP (green) signals. A reflection image (gold) was used to identify the lamella shape and location. **(B)** Cryo-CLEM image of lamella. Medium montage map image of lamella is aligned with and overlayed on cryo-confocal image from A. Black arrows indicate colocalization of fluorescent signals. **(C)** Zoomed images of areas indicated in B (i-iii). Cryo-confocal (left) and TEM image (right) are from the same location. Synaptic vesicle-containing compartments are well-aligned with presynaptic red fluorescence signal (asterisk). Scale bars are 500 nm.

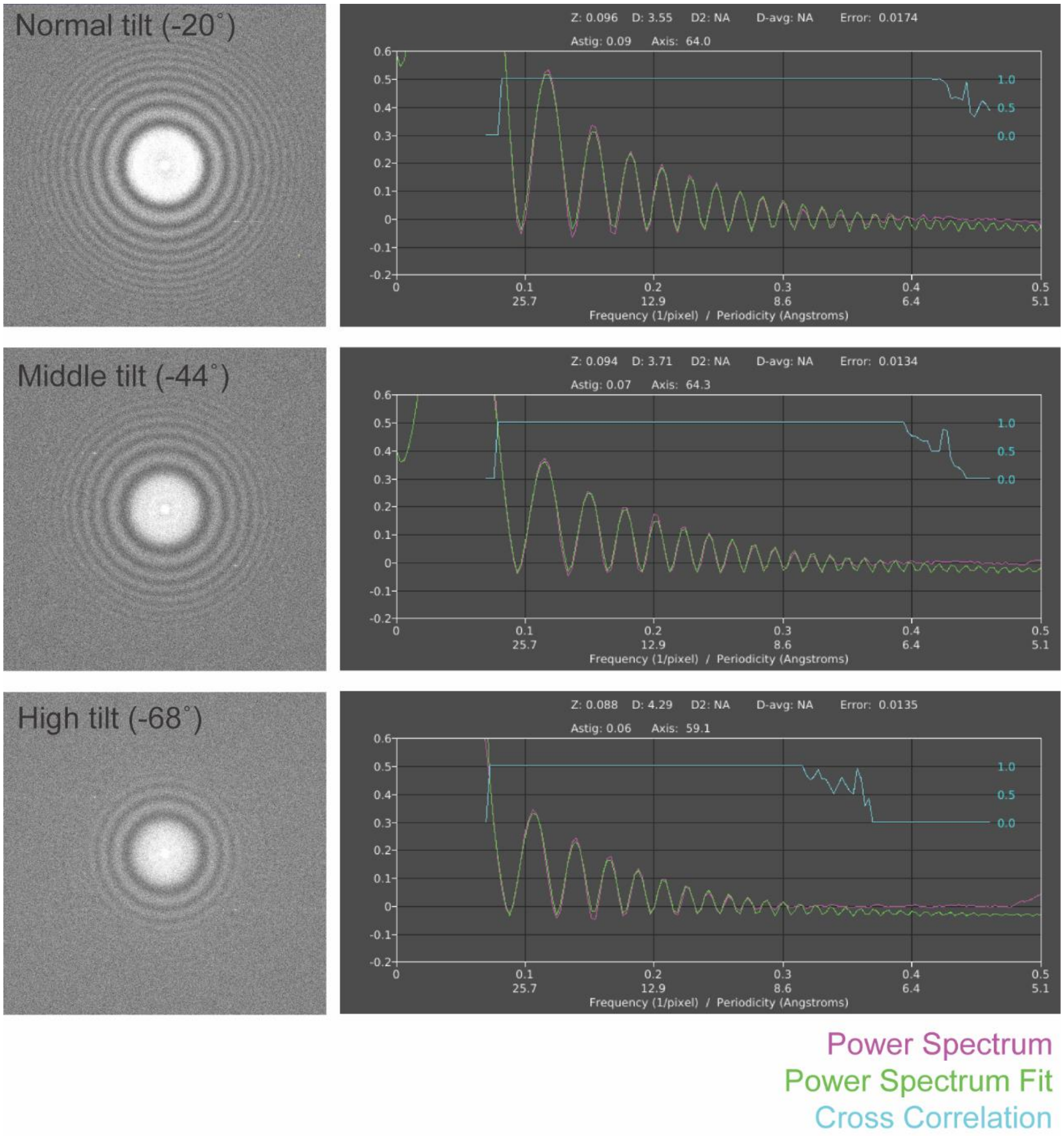


**Supplemental Figure 5. Power spectra and contrast transfer function (CTF) fitting of tilt series.** IMOD calculation of Fast Fourier Transform, or FFT (left) and Ctfplotter fits (right) for three tilts from a single representative tilt series.

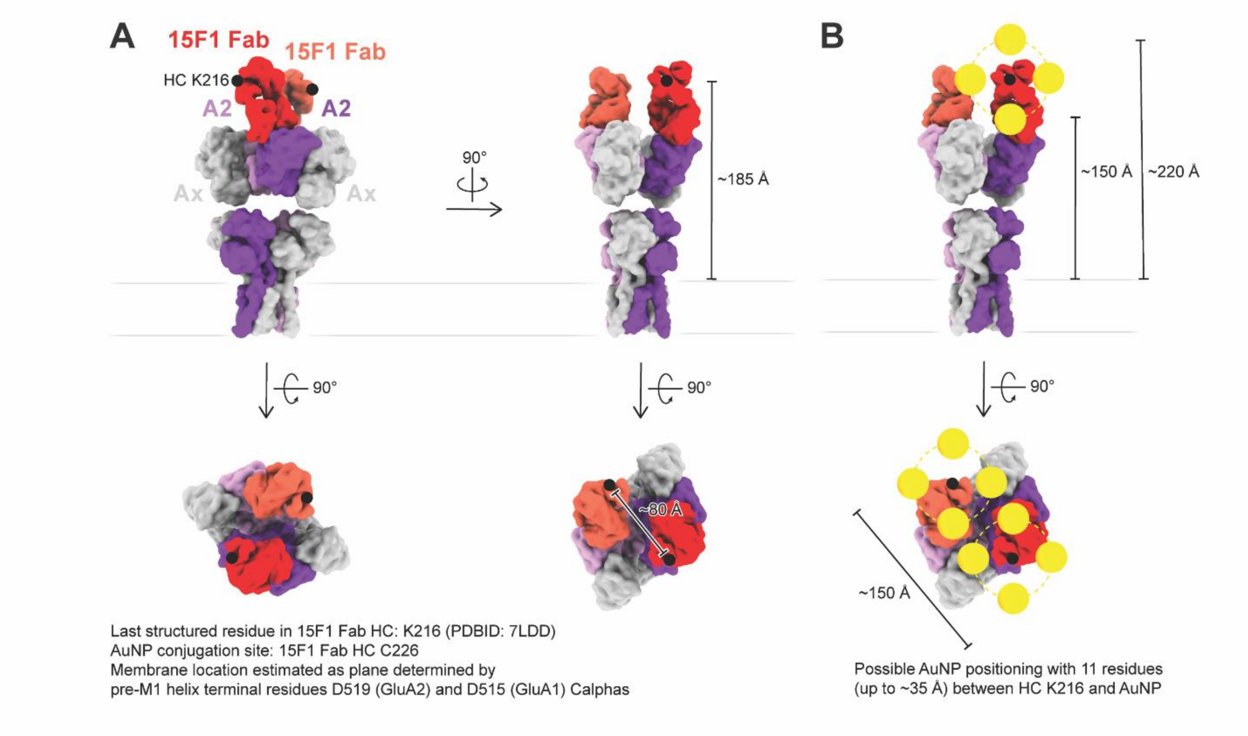


**Supplemental Figure 6. Analysis of spatial constraints of 15F1 Fab-AuNP conjugation. (A)** Structure of native hippocampal AMPAR bound to 15F1 Fab (PDB: 7LDD), with the position and relative distance between the last structured 15F1 Fab heavy chain residues, K216, on each Fab and between K216 and the membrane denoted. **(B)** Spatial representation of possible AuNP positioning in the 15F1Fab-AuNP:AMPAR complex, restraining the AuNP to a distance of 35 Å from K216, with ranges of inter-AuNP and AuNP-membrane distances denoted.

**Supplemental Video 1. Representative Tomogram and Segmentation.**

**(0:00-0:10)** Zoom from lamella overview to the area of interest. **(0:10-0:20)** Z-Slice animation of reconstructed tomogram (denoised using CryoCare). **(0:20-0:33)** Reverse Z-Slice animation revealing tomogram segmentation. **(0:33-0:54)** Pan around segmentation. **(0:54-1:14)** Close-up of synaptic cleft revealing organization of AuNPs.
